# Tracing endogenous proteins in living cells through electrotransfer of mRNA encoding chromobodies

**DOI:** 10.1101/2023.09.20.558581

**Authors:** Théo Juncker, Ludovic Richert, Murielle Masson, Bruno Chatton, Mariel Donzeau

## Abstract

Chromobodies made of nanobodies fused to fluorescent proteins are powerful tools for targeting and tracing intracellular proteins in living cells. Typically, this is achieved by transfecting plasmids encoding the chromobodies. However, an excess of unbound chromobody relative to the endogenous antigen can result in high background fluorescence in live cell imaging. Here, we overcome this problem by using mRNA encoding chromobodies. Our approach allows one to precisely control the amount of chromobody expressed inside the cell by adjusting the amount of transfected mRNA. To challenge our method, we evaluate three chromobodies targeting intracellular proteins of different abundance and cellular localization, namely lamin A/C, Dnmt1 and actin. We demonstrate that expression of chromobodies in living cells by transfection of tuned amounts of the corresponding mRNAs allows the accurate tracking of their cellular targets by time-lapse fluorescence microscopy.

## 1. Introduction

Real-time imaging of living cells is a very powerful and versatile tool to study the dynamics of biological processes.^[1]^ Ectopic expression of chosen cellular proteins, fused to appropriate fluorescent tags, is an invaluable tool for tracking a range of intracellular events on a single cell basis.^[1,2]^ For instance, key events of mitosis, such as chromosome segregation, have been studied using time-lapse fluorescence microscopy on mammalian cells stably expressing mCherry-tagged histone H2B.^[3]^ However, fluorescent tags directly fused to the protein of interest can interfere with specific localization signals, disrupt proper protein folding or oligomerization, or cause steric hindrance, preventing interactions with protein partners and subcellular structures.^[2]^ In addition, ectopic overexpression of tagged protein may not reflect the distribution or the dynamic process of the endogenous protein, as it has been observed for vimentin.^[4]^ It is therefore crucial to accurately identify and validate the spatial and temporal dynamics of endogenous proteins. To date, only few monoclonal antibodies (mAbs) or antibody fragments (Fab and scFv) have been successfully expressed or directly delivered as proteins to target intracellular antigens in living cells,^[5–11]^ but in most cases, their application has been hampered by inefficient folding and disulfide bond formation in the reducing environment of the cytosol, leading to aggregate formation in living cells.^[12]^

Recently, nanobodies or V_H_Hs (variable heavy chain of a heavy chain antibody) have emerged as valuable and powerful tools for tracking intracellular proteins, due to their small size (around 15 kDa), robustness and ability to fold in a reducing environment.^[13]^ V_H_Hs genetically fused to fluorescent proteins, called chromobodies, are now well established for monitoring the fate of endogenous proteins in living cells, by transient transfection of plasmids encoding chromobodies.^[13–15]^ However, transient expression of chromobodies displays a large intercellular heterogeneity and produces an excess of V_H_H versus the endogenous antigen,^[14,15]^ so that the generation of cell lines stably expressing low levels of chromobody is generally more reliable. Stable cell lines expressing an actin-chromobody and a vimentin-chromobody were successfully generated to follow the spatiotemporal dynamics of both endogenous proteins using live cell imaging.^[14,15]^ Another approach to avoid an excess of nanobody over endogenous antigen is to transduce purified nanobodies into living cells. This technique was successfully applied for the visualization of H2AX phosphorylation at the Serine 139, specifically γ-H2AX, through the direct delivery of the purified chromobody C6B-tagged-dTomato.^[16]^ Nevertheless, this process required production and purification of the chromobody from *E*.*coli*, which may be laborious and time consuming.

Recently, we successfully detected endogenous proteins using electrotransfer of *in vitro* transcribed (IVT) mRNAs encoding chromobodies.^[17]^ We have shown that mRNA is ideal for delivering genetic information into various cell lines with extremely high efficiency, up to 95%. Furthermore, by monitoring the amount of transfected mRNA, we were able to adjust the amount of nanobody expressed to the amount of target antigen inside the cell, eliminating the need to remove the excess of unbound nanobody.^[17]^ In the present work, we describe the huge potential of mRNA encoding chromobody delivery for detecting and tracing intracellular targets with extremely high accuracy by live cell imaging. As a proof of concept, we tested three chromobodies to visualize the dynamic behavior of endogenous proteins present in different abundance and in distinct subcellular compartments, by real-time microscopy; the lamin A/C components of the nuclear lamina^[13]^, the β-actin, highly conserved protein and involved in various types of cell mobility^[18]^ and the DNA methyltransferase 1 (Dnmt1) that methylates CpG residues and is associated with DNA replication sites during in S phase.^[19]^ Because the mRNAs are directly delivered into the cytoplasm, expression occurs in almost all cells as early as 3 hours after transfection without cell division and lasts for at least for 48 h. In principle, this approach is not limited to chromobodies, but can be applied to any other specific intracellular protein binders.

## 2. Results and Discussion

### 2.1. Lamin-chromobody

We first evaluated the anti-nuclear lamin VHH binding to lamin A/C.^[13]^ The nanobody coding sequence was cloned in frame with either the mScarlet,^[20]^ the mScarlet fused to E3 peptide originated from the tetramerization domain of p53 ^[17]^ or the dTomato coding sequences^[21]^, the last two both forming dimers (Figure 1A). The DNA constructs were then used for IVT mRNA synthesis. Different amounts of mRNAs coding for the three chromobodies were electrotransfected into the HeLa H2B-GFP cell line. The endogenous lamina was clearly detected by all three chromobodies in almost 95% of the cells, although with different efficiencies (Figure 1B). Both dimeric chromobodies gave a strong and specific signal at 100 ng of transfected mRNA, while the signal of the monomeric lamin-chromobody tagged mScarlet was very weak at 200 ng and partly unspecific. The fact that both dimer formats were significantly better at detecting the nuclear lamina than the monomeric lamin-chromobody resulting in a higher signal-to-background ratio is not surprising. Indeed, bivalent V_H_Hs have been shown to display a better functional affinity compared to their monovalent counterparts through avidity effect.^[16,22]^ However, the lamin-chromobody tagged dTomato gave a stronger signal than the lamin-chromobody mScarlet-E3 with equivalent amounts of transfected mRNA (figure 2B). This observation was surprising as the mScarlet and dTomato have an equivalent brightness signals *in cellula*.^[23]^ Nevertheless, aggregates were clearly detected for the lamina-chromobody tagged dTomato (Figure 1B and C). This phenomenon was not observed with the lamina-chromobody tagged mScarlet-E3. Therefore, we decided to use with lamina-chromobody tagged mScarlet-E3 construct.

**Figure 1:**
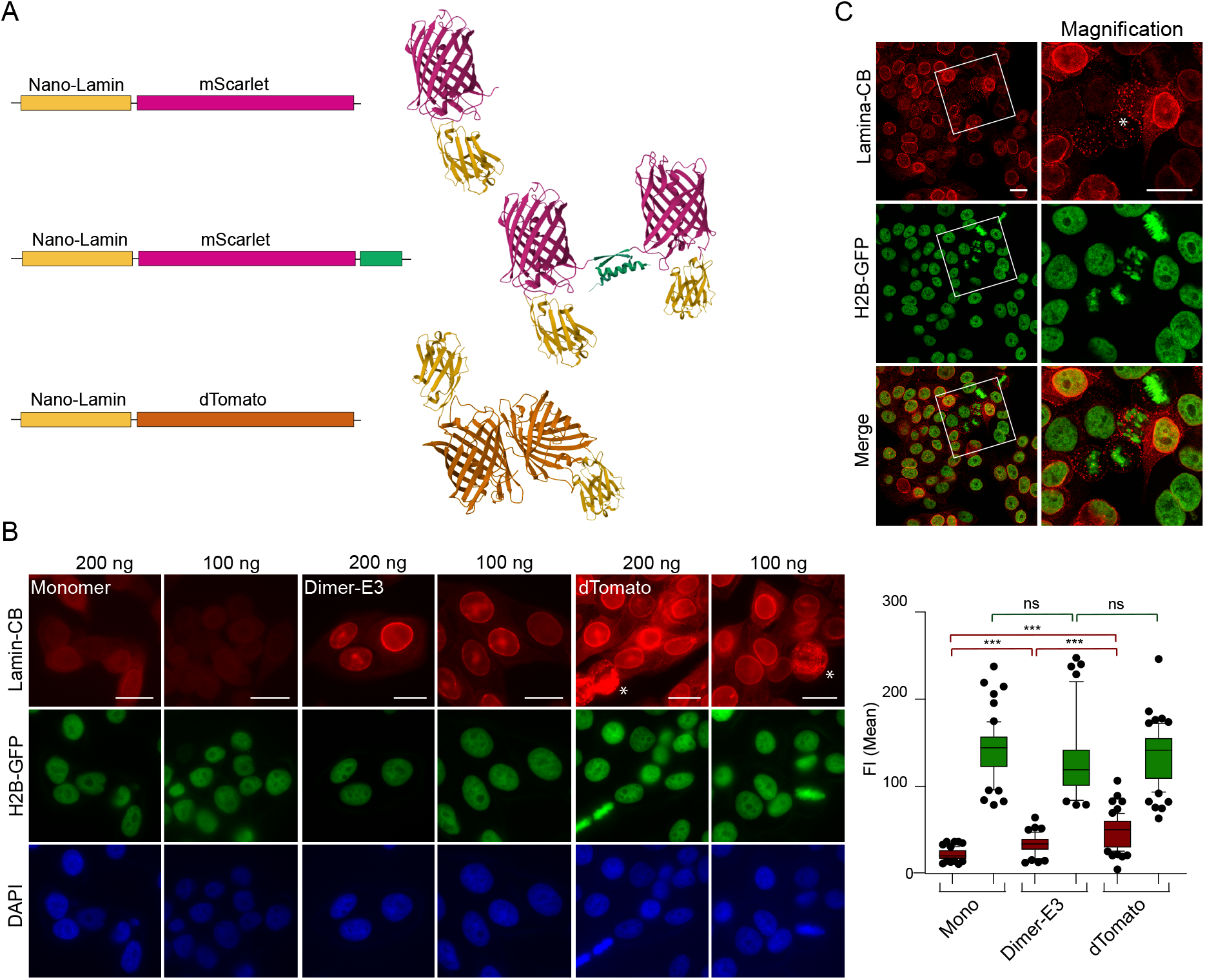
Determination of the optimal amount of transfected mRNAs for different lamin-chromobodies (Lamin-CB). (A) Schematic representations of the different Lamin-chromobody constructs; in the left panel; yellow box for nanobody anti-lamin (nano-Lamin), purple box for the mScarlet, orange box for the dTomato and green box for the E3 helix. The oligomerization states of the corresponding encoded proteins are depicted on the right panel. (B) HeLa-H2B-GFP cells were electrotransfected with various amounts of mRNAs encoding lamin-chromobody (Lamin-CB) constructs tagged mScarlet (Monomer), lamin-chromobody tagged mScarlet-E3 (Dimer-E3) and lamin-chromobody tagged dTomato (dTomato), as indicated. Images were taken using fluorescent microscope Leica. The quantification of the mean fluorescence intensity (FI) of cells transfected with the three chromobodies are represented, with in red the FI mean of the mScarlet signal and in green the FI mean of the GFP signal. The Student’s *t*-test was performed: ns: not significant, \*\*\**p*-value<0.0001. (C) HeLa-H2B-GFP cells were electroporated with 100 ng of mRNAs encoding lamin-chromobody tagged dTomato. Aggregates are highlighted with an asterisk (*). Scale bar, 20 μm.

**Figure 2:**
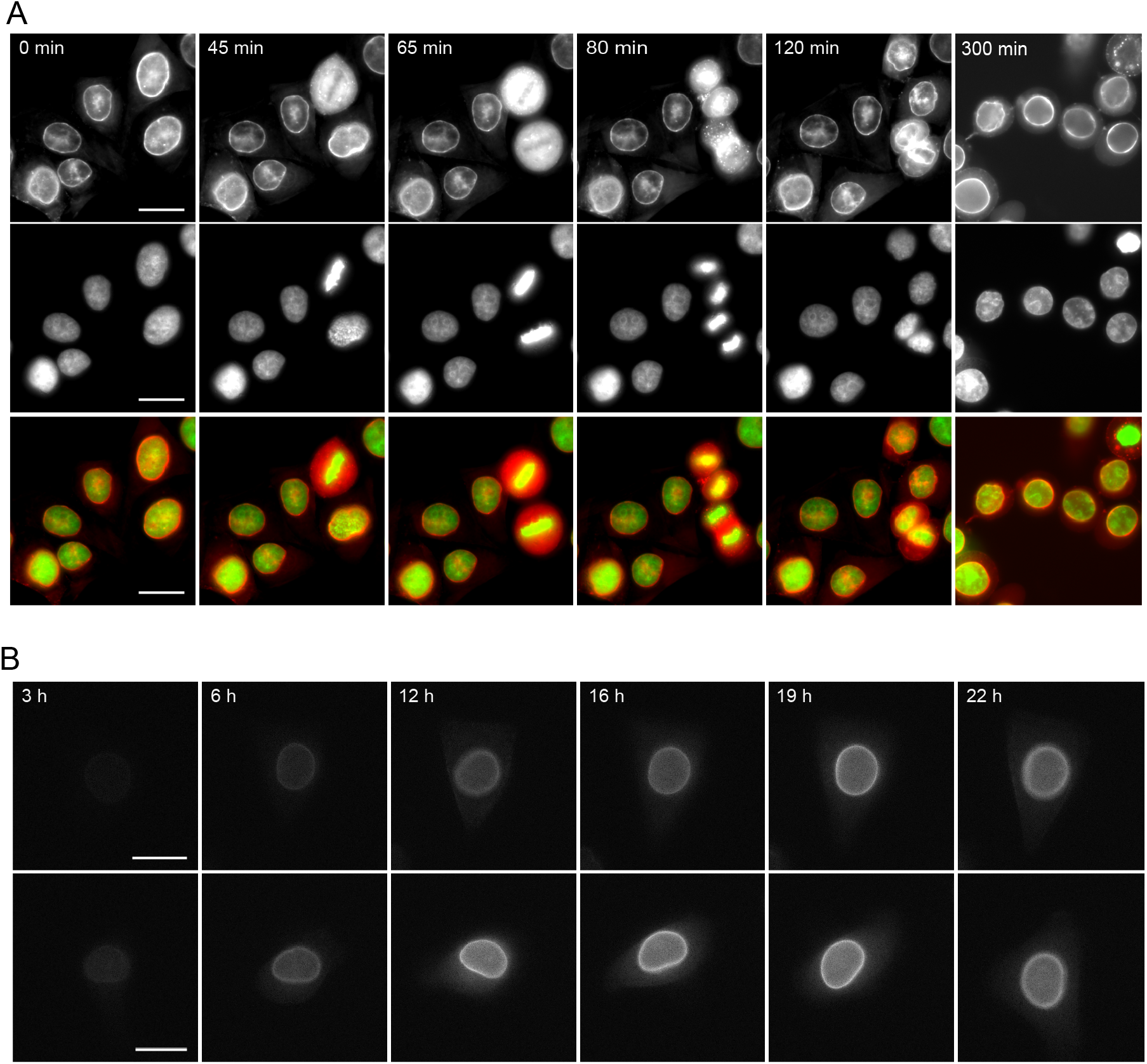
Live-cell imaging of HeLa-H2B-GFP cell-line transfected with the mRNA encoding lamin-chromobody mScarlet-E3 starting either (A) forty-eight hours or (B) three hours after transfection. (A) The red signal of lamina-chromobody mScarlet-E3 is merged with that of the H2B-GFP. Bleach correction has been applied using histogram matching in Image J. (B) The lamina-chromobody is depicted in grey. In order to monitor the increasing signal, no bleach correction was applied to show the increase in lamin-chromobody expression. Scale bar 20 μm.

Thus, 100 ng of lamin-chromobody mScarlet-E3 encoding mRNA was electroporated in HeLa-H2B-GFP cells. Forty-eight hours later, the dynamics of the lamina was observed at different stages of the cell cycle (Figure 2A). In G1 and S phases, the lamin-chromobody mScarlet-E3 signal was specifically distributed at the periphery of the nucleus. During the mitosis, it was homogeneously distributed throughout the cytoplasm of the dividing cells (Figure 2A and Movie 1). As the mRNA is delivered in the cytoplasm, we tested whether the IVT mRNA was rapidly translated. A time point expression of the lamin-chromobody was performed following transfection of 100 ng of mRNA. As shown in figure 2B and Movie 2, the lamin-chromobody expression took place 4 h after mRNA delivery and peaked at around 16 hours after transfection (two cells are shown in figure 2B).

### 2.3. Dnmt1-chromobody

We next evaluated the nanobody targeting the human DNA methyltransferase 1 (hDnmt1).^[24,25]^ In the nucleus, Dnmt1 associates with the replication factories, a mechanism that links the maintenance of the genomic methylation patterns to the process of DNA replication.^[26,27]^ Dmnt1 transfers methyl groups to hemimethylated substrate CpG sites generated during DNA synthesis in S phase.^[28]^ In early and mid S phase, Dnmt1 transiently interacts with the proliferating cell nuclear antigen (PCNA), whereas in late S phase, Dnmt1 concentrates at chromocenters, which are composed of the pericentric heterochromatin of individual chromosomes.^[27,29,30]^ Consequently, the distribution pattern of Dnmt1 is highly variable throughout the cell cycle.

The nanobody anti-Dmnt1 coding sequence was cloned in frame with the mScarlet and the E3 peptide to obtain a dimer (Figure 3A) and the DNA construct was used for mRNA IVT. Then, various amounts of mRNA encoding the Dnmt1-chromobody mScarlet-E3 were electro-transfered into two different cell lines, MRC5 and HeLa-H2B-GFP. Twenty-four hours after transfection, the cells were fixed and observed by direct immunofluorescence. At levels of 200 and 100 ng transfected mRNA, the Dnmt1-chromobody mScarlet-E3 was diffusely present in all compartments of the cell, whereas at 50 ng the Dnmt1 signal was predominantly localized in the nucleus in both cell lines (Figure 3B). To obtain a specific signal with minimal background, the optimal quantity of mRNA was 50 ng for HeLa H2B-GFP and 25 ng for the MRC5 (Figure 3B and C and Figure S1). Importantly, with 50 ng of mRNA transfected, almost 95% of the cells were expressing the Dnmt1-chromobody mScarlet-E3 (Figure 3B and C, and Figure S2).

**Figure 3:**
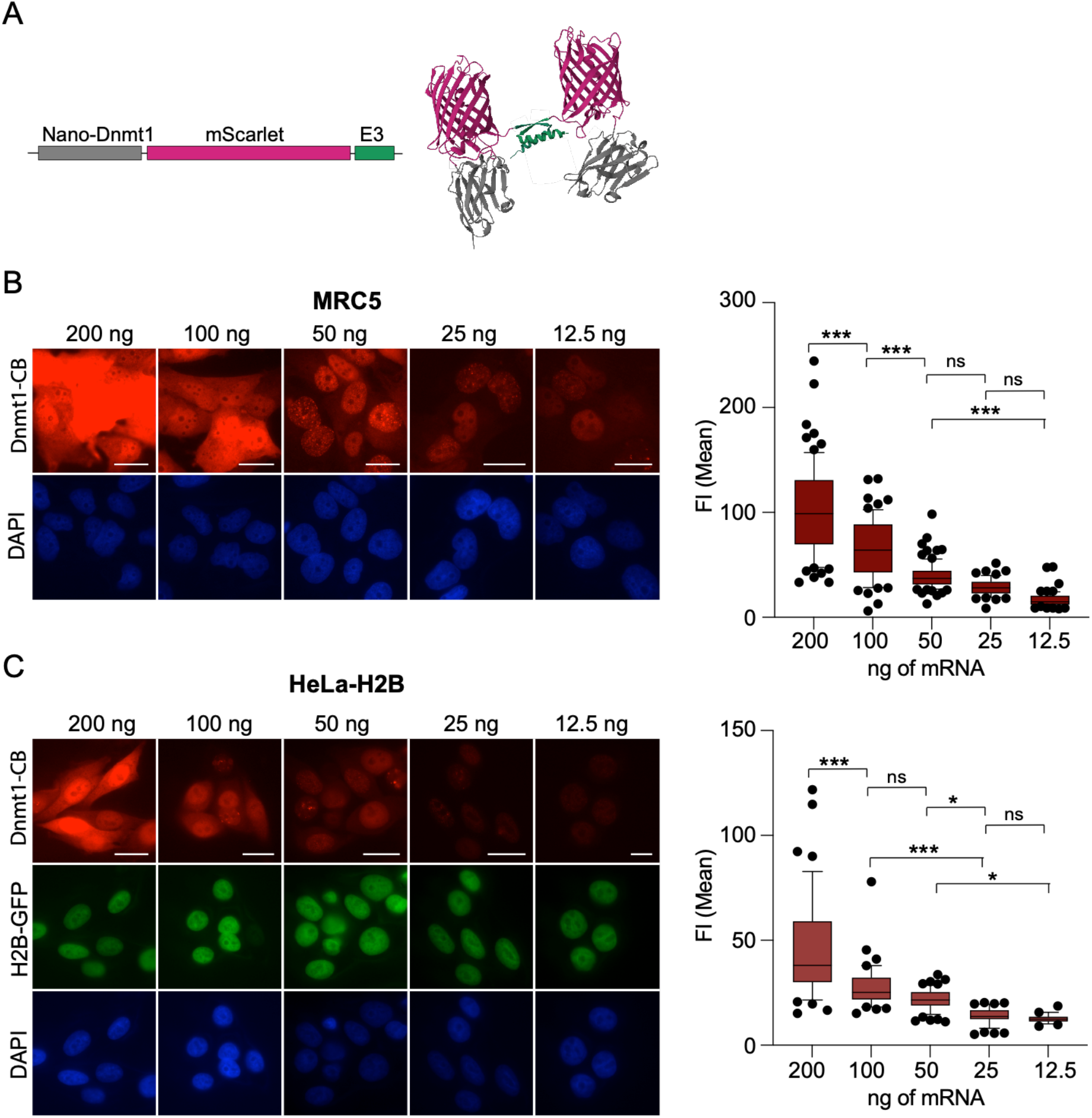
Estimation of the amount of mRNA to be electrotransfected for an optimal detection of the Dnmt1 protein with a minimum ratio of signal to background. Various amounts of mRNA encoding - Dnmt1-mScarlet-E3 chromobody were transfected either (A) in MRC5 or (B) in HeLa-H2B-GFP cell-lines, as indicated. Twenty-four hours after transfection, cells were fixed and images with an identical exposure time were taken for analysis. The quantification of the mean fluorescence intensity (FI) of cells transfected with the different amount of mRNAs encoding Dnmt1-chromobody is represented. Scale bar, 20 μm.

Next, MRC5 cells electrotransfected with the optimal amount of mRNA, were monitored for Dnmt1 dynamics distribution using live-cell imaging. As shown previously,^[24,29]^ the signals are characteristic of Dnmt1 dynamics during the cell cycle, corresponding to the spatial and temporal coupling of Dnmt1 with sites of active DNA replication during early and late S-phase, when DNA of percentromeric heterochromatin is replicated (Figure 4A, Movie 3). The co-localization of Dnmt1 and PCNA during S-phase was demonstrated by immunofluorescence on fixed cells expressing Dnmt1-chromobody using a mAb anti-PCNA (PC10) (Figure 4B). Thus, in S-phase, Dnmt1 transiently co-localizes with PCNA foci and then concentrates at chromocenters.

**Figure 4:**
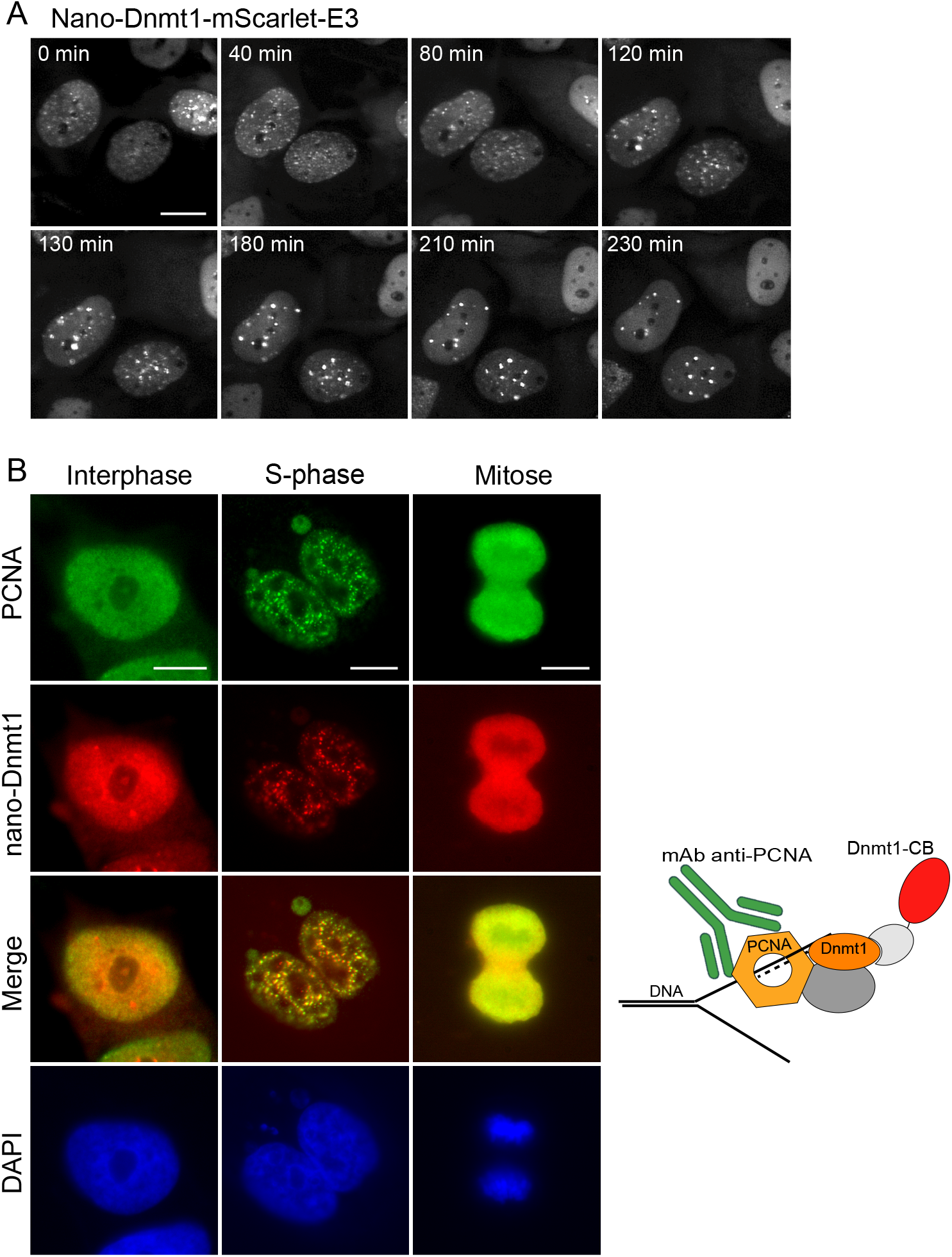
Dnmt1 dynamics in living cells. Live cell imaging of (A and B) MRC5 cells transfected with 50 ng of mRNA encoding the Dnmt1-mScarlet chromobody. (A) Twenty-four hours after transfection, cells were analyzed by live cell imaging. Individual time frames reveal endogenous Dnmt1 distribution at the indicated time points as assessed in Movie 3. Scale bar, 20 μm. (B) Co-localization of Dnmt1 and PCNA during the cell cycle. 24 h after mRNA transfection, cells were fixed and probed for PCNA using a mouse-mAb anti-PCNA (PC10) and a secondary anti-mouse Alexa488. Scale bar, 10 μm.

We evaluated the expression level of the Dnmt1-chromobody at different time points with either 50, 100 or 1000 ng of transfected mRNA in MRC5 cell line. The Dnmt1-chromobody was expressed as early as 3 h post-transfection, respectively (Figure 5A and Figure S3). The fluorescence intensity was clearly proportional to the amount of transfected mRNA and the time point measurement after transfection. Interestingly, 20 h after transfection, the majority of the Dnmt1-chromobody signal was localized in the nucleus, whereas 6 h after transfection Dnmt1-chromobody signal was detected in the nucleus and the cytoplasm. The dimeric Dnmt1-chromobody with a size of around 100 kDa can not diffused freely through the nuclear pore complex, and since it lacks a nuclear localization signal (NLS), Dnmt1-chromobody is likely to enter the nucleus via a piggybacking mechanism.^[31,32]^ Indeed, Dnmt1-chromobody might piggyback indirectly via its physical interaction to *de novo* synthetized Dnmt1 protein encompassing three NLS that directly binds the nuclear transport apparatus.^[33]^ Thus, we hypothesized that the mRNA might be degraded rapidly and the produced Dnmt1-chromobody excess, which has accumulated in the cytoplasm, is then gradually transported in the nucleus by binding to the newly synthetized Dnmt1 protein. The amount of the mRNA encoding Dnmt1-chromobody was quantified by reverse transcriptase quantitative PCR (RT-qPCR) assay in electrotransfected mRC5 cells. The level of mRNA encoding Dnmt1-chromobody peaked at 2 hours post-transfection, decreased rapidly at 6 hours and was nearly undetectable at 20 hours post-transfection (Figure 5B). It then decreased two-fold 3 hours later and became undetectable 20 hours after transfection. This observation supports our hypothesis that the translated mRNA leads to some accumulation of the Dnmt1-chromobody in the cytoplasm and then, as the mRNA is degraded, the Dnmt1-chromobody is gradually transported to the nucleus by binding to newly synthesized Dnmt1 (Figure 5C).

**Figure 5:**
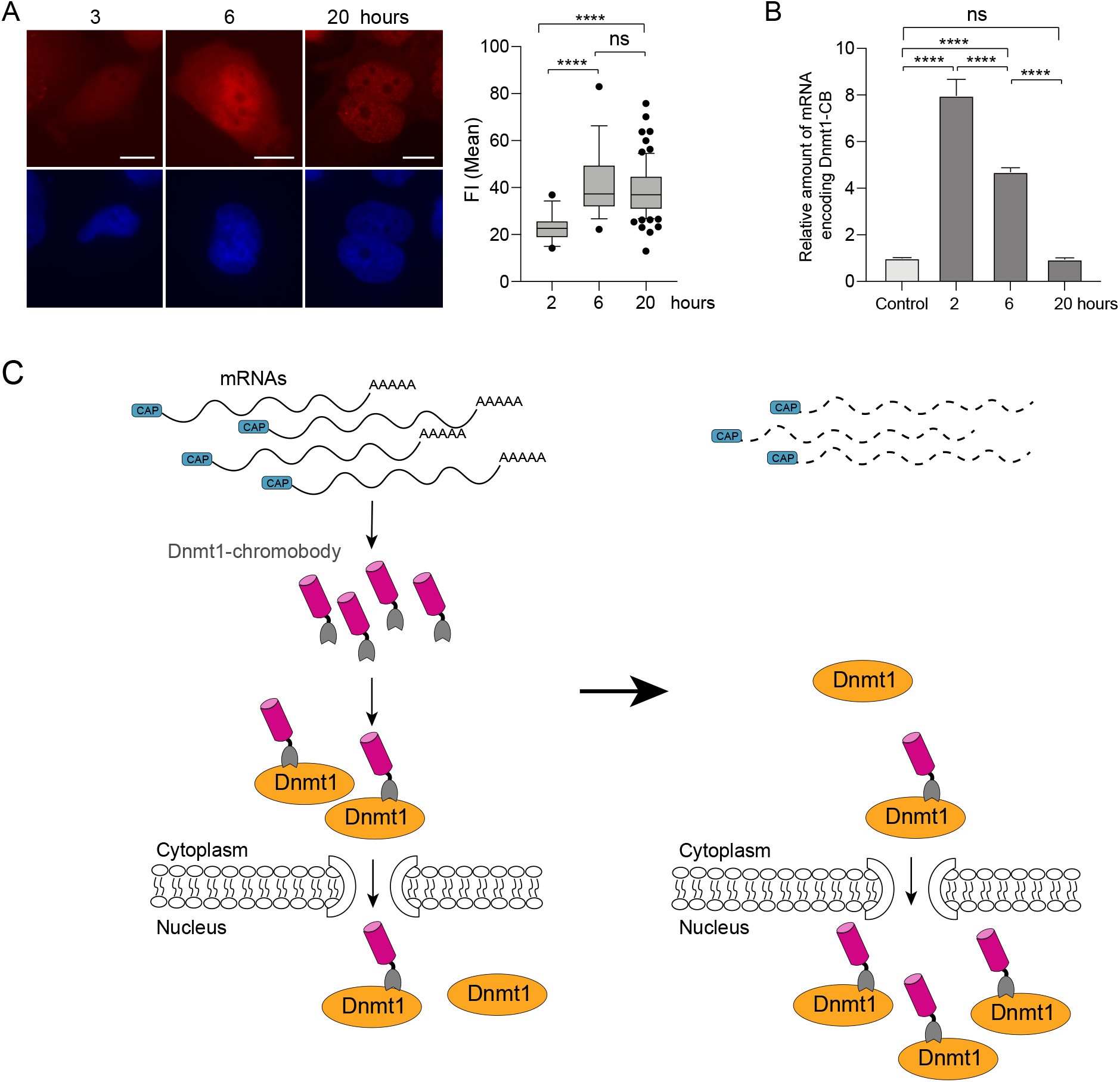
Correlation between mRNA levels and with Dnmt1-chromobody expression in the MRC5 cell line. Cells were transfected with 50 ng of Dnmt1-chromobody encoding mRNA. Cells were fixed at different times after transfection, and images taken at the same exposure time for analysis (A). Mean fluorescence intensity (FI) quantification is shown. (B), Cells were harvested for qPCR analysis to determine the relative levels of Dnmt1-CB mRNAs. (C) Schematic representation of the piggyback mechanism for the Dnmt1-chromobody bound to *de novo* Dnmt1 protein.

### 2.4. Actin-chromobody

Finally, we evaluate the actin-chromobody tagged RFP. Actins represent the most abundant proteins present in most eukaryotic cells, which participate in multiple important cellular processes including cell mobility, cell division and organelle dynamics.^[18]^ First, the amount of mRNA to be transfected was determined. A specific and accurate signal of the actin network was detected when 200 ng of mRNAs encoding actin-chromobody were transfected in MRC5 cells (Figure 6A). Live cell imaging was performed on MRC5 and U2OS cell lines transfected with 200 ng of mRNAs encoding actin-chromobody (Figure 6 B-C and Movies 4 and 5). Clear detection of actin filaments during cell division or cell mobility were observed by live-cell imaging validating the transfection of mRNA encoding actin-chromobody to trace actin dynamic in living cell.

**Figure 6.**
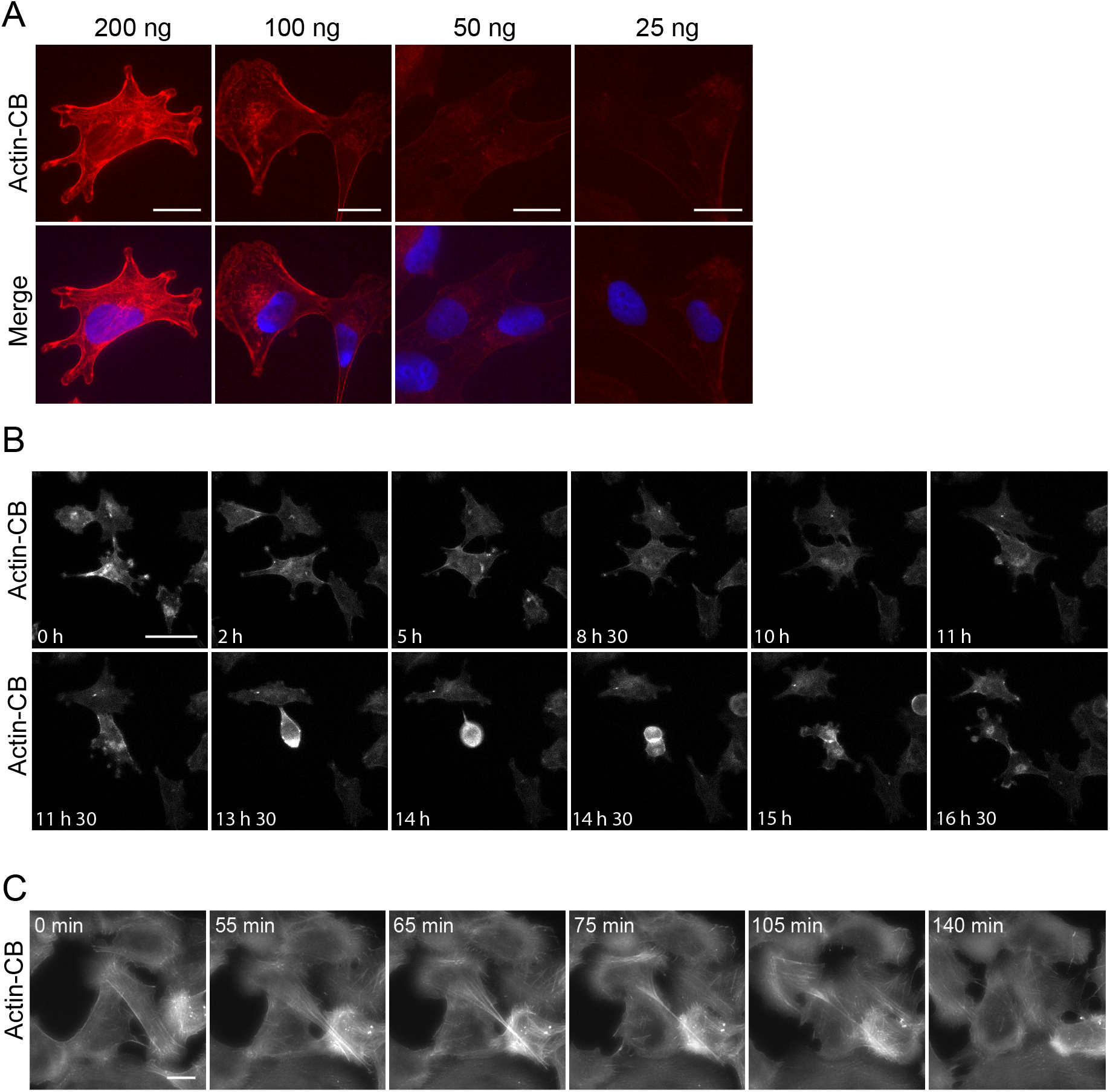
Actin visualization using transfection of mRNA encoding actin-chromobody. (A) Estimation of the optimal amount of mRNA encoding actin-chromobody to be transfected in MRC5. Different amounts of mRNA were transfected in MRC5, as indicated. Cells were fixed and analyzed by fluorescence microscopy. (B and C) 200 ng of mRNA encoding actin-chromobody were transfected in MRC5 cells (B) or U20S (C). Twenty-four hours after transfection, cells were analyzed by live cell imaging. Individual time frames reveal endogenous actin at the indicated time points as assessed in Movie 4 and 5. Scale bar, 20 μm.

## 3. Conclusion

Chromobodies offer a tremendous advantage for the direct detection and visualization of almost any endogenous protein within living cells by real-time microscopy. Transient transfection of plasmid encoding the chromobodies has been used.^[14–16]^ However, transfection of expression plasmids almost always results in an excess of nanobody compared to the native antigen. For instance, the spatiotemporal dynamic changes of the endogenous vimentin have been studied by time-lapse imaging using transient transfection of a vimentin chromobody.^[15]^ However, as the transient chromobody expression showed a large intercellular heterogeneity, a stable chromobody cell line was established and selected for low chromobody expression to minimize background fluorescence due to an excess of unbound chromobody.^[15]^ Likewise, the endogenous nuclear actin was visualized using the actin-Chromobody targeted to the nucleus.^[14]^ Stable nuclear actin-chromobody expressing cells were generated to better monitor nuclear actin assembly more accurately than with transient transfection.^[14]^ The actin-chromobody was also used to visualize the sub-organelle actin spatiotemporal dynamics with high resolution,^[34]^ but over-expression artifacts were observed due to the strong CMV promoter driven transient expression of the nanobody. To overcome this problem, the authors expressed the actin-chromobody using a weaker promoter.^[34]^ A second strategy consists of directly transducing purified chromobodies as proteins in living cells.^[16]^ In such case, the levels of nanobodies are generally lower than those of the target antigen. Nonetheless, aside from being a laborious and time consuming methodology, expression levels, folding and solubility of the chromobody may turn out to be challenging.^[16]^

Here, we establish a simple and straightforward alternative to detect and track endogenous proteins in real time using mRNAs encoding chromobodies. The mRNAs are synthetized *in vitro* and delivered into living cells by electroporation.^[17]^ The possibility to precisely control the level of chromobody expressed by adjusting the amount of mRNAs transfected ensures the ability to fine-tune the level of chromobody with its cellular target. This was demonstrated by targeting three endogenous proteins of different abundance, namely β-actin, lamin A/C and Dnmt1. Indeed, the β-actin is approximately 5-fold more abundant than lamin A/C, which is itself 30-fold more abundant than Dnmt1.^[35,36]^

We strongly believe that the delivery of mRNA encoding chromobodies opens up many possibilities for the specific generation of probes to accurately unravel the spatio-temporal distribution of intracellular components in living cells.

## Materials and Methods

### Cell lines

HeLa-H2B-eGFP and MRC5 (ATCC CCL-171) were maintained as monolayers in Dulbecco’s modified Eagle’s medium (DMEM) (1 g/L glucose) (Gibco, Billings, MT, USA) supplemented with 10% heat-inactivated FBS and Pen–Strep.

### Plasmids

The coding sequence of the Dnmt1-chromobody targeting the human Dnmt1 protein ^[37]^ (Uniprot : P26358 · DNMT1_HUMAN) and lamin-chromobody (ChromoTek) targeting the lamin D0 from Drosophila monogaster (Uniprot : P08928 · LAM0_DROME), were amplified by PCR with primers NaBo Foward CGTCAGCCATGGGGTCCCAGGTTCAGC and NaBo Reverse-STOP CCACAGGAATTCACAATGGTGATGATGGTGATGTGCG and subcloned into the NcoI-EcoRI digested pETOM to obtain the pETOM-nano-Dnmt1 and the pETOM-nano-Lamina.^[38]^ The mScarlet-E3 was amplified from the pET-mScarlet-E3 described elsewhere^[39]^ with primers mScarlet-SpeI-For GGGCCCACTAGTGTGAGCAAGGGCGAGGCAGTGATC and E3/K3-EcoRI-Rev CATGACGAATTCTTAACCCCCTGGCTCTTCCC, and subcloned into SpeI-EcoRI restriction sites of the pETOM-nano-Lamin giving rise to the pETOM-nano-Lamin-mScarlet-E3. The dTomato DNA sequence was amplified from pET-C6B-dTomato with primer dTo-For-SpeI CATGATACTAGTATGGTGAGCAAGGGCGAGGAGG and a reverse primer dTo-Rev-EcoRI GTGCTCGAATTCTTATTAATGG, and subcloned into the SpeI-EcoRI digested pETOM-nano-Lamina to obtain the pETOM-nano-Lamin-dTo. The plasmid encoding the actin-chromobody (ChromoTek) was used to amplify the coding sequence of the chromobody under the control of the T7 promoter with actin-chromobody-for AATTAATACGACTCACTATAGGGAGAAAGGAGATATCCATGGCTCAGGTGCAGCTGGTGG and chromobody-RFP/GFP-rev TTATGATCTAGAGTCGCGGCCGC.

### mRNA synthesis and transfection

The mRNAs were transcribed from the T7 promoter of a linearized pETOM vector with EcoRI restriction enzyme containing the gene of interest. Linearized plasmids were purified and used as a template for in vitro transcription (IVT) as previously described.^[17]^ The transfection experiments with mRNAs were conducted with the NeonTM transfection system (Thermofisher Scientific, Waltham, MA, USA) as previously described.^[17]^ The cells were transferred into 24-well plates containing glass coverslips or in Ibidi for time-lapse imaging.

### Immunofluorescence

The cells grown on coverslips were fixed with 4% (*w*/*v*) paraformaldehyde for 20 min. Coverslips were mounted with Fluoromount G containing 4′,6′-diamidino-2 phenyleindole (SouthernBiotech, Birmingham, UK) and cells were observed by direct immunofluorescence. For indirect fluorescence, cells were prior incubated with 100% cold-methanol for 5 min prior to be incubate with an anti-PCNA mouse mAb (PC10) in PBS containing 0.2 % Triton X100 and 6% FCS followed with a secondary anti-mouse-Alexa 488 (Life Technology, Carlsbad, CA, USA). The treated cells were analyzed using conventional fluorescence microscopy with a Leica DM5500 microscope (Leica Microsystems, Wetzlar, Germany). For time-lapse, Images were processed with Image J2.0.0.

### Time-lapse microscopy

Time-lapse movies were recorded using inverted microscopes either DMIRE2, Leica, equipped with a 40X NA 0.75 objective (HCX PL APO, Leica), a solid-state light source (SOLA III, Lumencor), a sCMOS camera (Prime, Photometrics) or Olympus IX83 with in both cases an environmental control system (Life Imaging Service). Cells were grown on μ-Dish 35mm (Ibidi) and maintained at 37°C in a 5% CO2 humidified atmosphere. Images were acquired at different intervals, depending on the experiment procedure for a maximum of 24 h using Metamorph (7.8.13.0, Molecular Device) or Olympus camera. Acquisition parameters were chosen to optimize signal to noise ratio and were fixed throughout the time-lapse acquisition. The resulting image stacks were processed using Fiji (ImageJ, NIH). No filtering procedure was used to visualize the dynamics of the lamina, the actin and the movement of Dnmt1 during the cell cycle.

### qRT PCR

Electrotransfected MRC5 cells were harvested and total mRNAs were purified with Nucleo Spin RNA kit (Macherey Nagel) according to the manufacturer’s instructions. cDNA was generated from 1 μg of RNA with High-capacity cDNA Reverse Transfection kit (Applied Biosystems). Real-time qPCR was performed with Syber Green I Master kit (Roche) on a Light Cycler LC480 instrument (Roche). For the human GAPDH gene, forward primer 5’-GCACAAGAGGAAGAGAGAGACC and reverse primer 5’-AGGGGAGATTCAGTGTGGTG were used. For the Dnmt1-chromobody, the forward primer 5’-CAGCGGTAACAACACCAGCTCC and reverse primer 5’-GGCATCCTCGAGTTCCAAGGCC were used to amplify a sequence within E3 sequence (from the tetramerization domain of p53). The Dnmt1 chromobody mRNA levels were normalized to the GAPDH mRNA levels. Changes in mRNA level were calculated by the threshold cycle (2^-ΔΔCT^) method. A one-way ANOVA tailed Student’s t test was performed on Δ*CT* values to determine the *p*-value.

### Statistics

The quantification was performed using Image J, and the statistic analysis was performed using GraphPad Prism 9.0. The number of replicates is indicated in the figure legends. In the boxplots, the bars indicate the median and interquartile range of the recorded fluorescence. The statistical significance of the data was determined with one-way ANOVA tailed Student’s t test and indicated as ns; not significant, * *p*-value <0.1, ** *p*-value <0.01, *** *p*-value <0.001 and **** *p*-value < 0.0001.

## Supplementary Materials

### Author Contributions

“Conceptualization, M.D.; methodology, M.D.; validation M.D.; investigation, T.J, L.R, M.M and M.D.; writing—original draft preparation, M.D.; writing— review and editing, M.M, B.C. and M.D.; supervision, M.D.; project administration, B.C.; funding acquisition, M.D. All authors have read and agreed to the published version of the manuscript.”

### Funding

This work was supported by funds and/or fellowships from the Centre National de la Recherche Scientifique, the University of Strasbourg, the French Ministry of Research, the Ligue Régionale contre le Cancer (CCIR-GE), the Fondation ARC pour la recherche sur le cancer, the ITI Innovec (IdEx) (ANR-10-IDEX-0002) and the SFRI (ANR-20-SFRI-0012).

## Acknowledgments

We thank G. Travé and K. Zanier for very helpful advises, J.C. Amé for the time-lapse microscopy on Olympus IX83, and the imaging was supported by the Imaging Center PIQ-QuESt (https://piq.unistra.fr/).

## Conflicts of Interest

The authors declare no conflict of interest.

## Notes

### Competing Interest Statement

The authors have declared no competing interest.

