## Supplemental figures for "Tracing endogenous proteins in living cells through electrotransfer of mRNA encoding chromobodies"

1 Biotechnologie et Signalisation Cellulaire (BSC), UMR7242, Université de Strasbourg, 300 Bvd Sebastien Brant, F-67412 Illkirch, France
2 Laboratoire de Biophotonique et Pharmacologie, (LBP) UMR 7213 CNRS, Université de Strasbourg, Faculté de pharmacie, 74 Route du Rhin, F-67401 Illkirch, France
3
4 Institut de Genetique et de Biologie Moleculaire et Cellulaire (IGBMC), INSERM U1258/CNRS UMR 7104/Université de Strasbourg, 1 rue Laurent Fries, BP 10142, F-67404 Illkirch, France

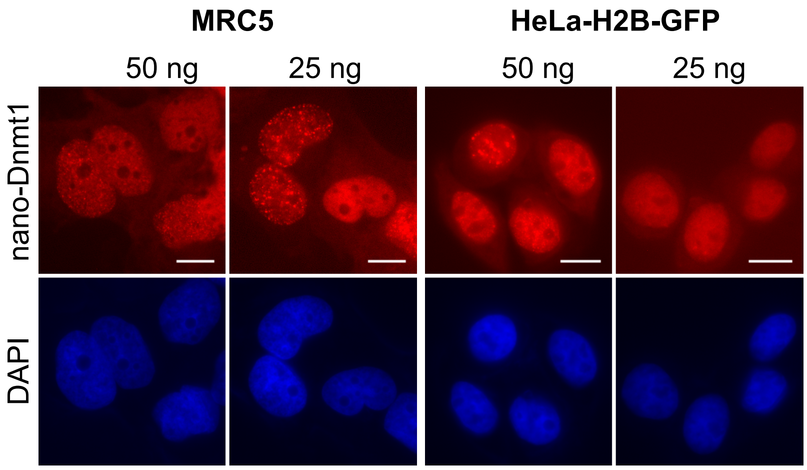


**Figure S1**: Images from the figure 3B and C were taken with different exposures to visualize the specificity of the Dnmt1-chromobody signals.

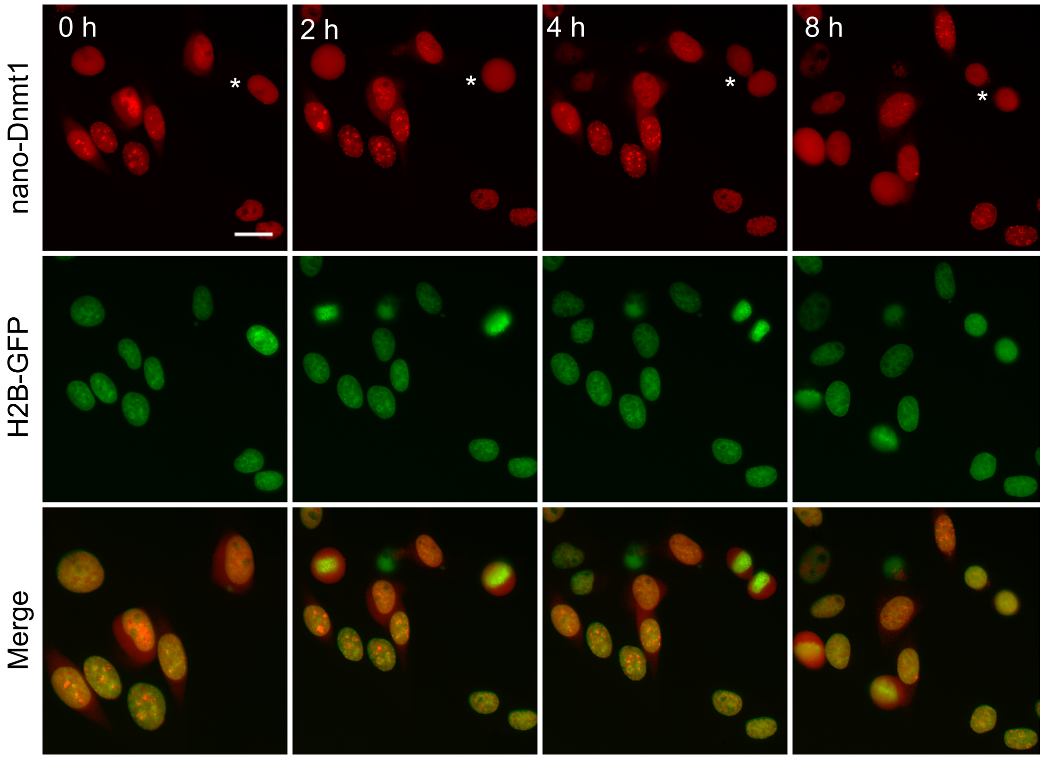


**Figure S2:** Expression of Dnmt1 chromobody in HeLa-H2B-GFP cell line. Time lapse was done 24 h after electroporation of 50 ng of mRNA encoding Dnmt1 chromobody. The signal of the chromobody (red signal) is merged with that of the H2B-GFP (green). Scale bar, 40 µm.

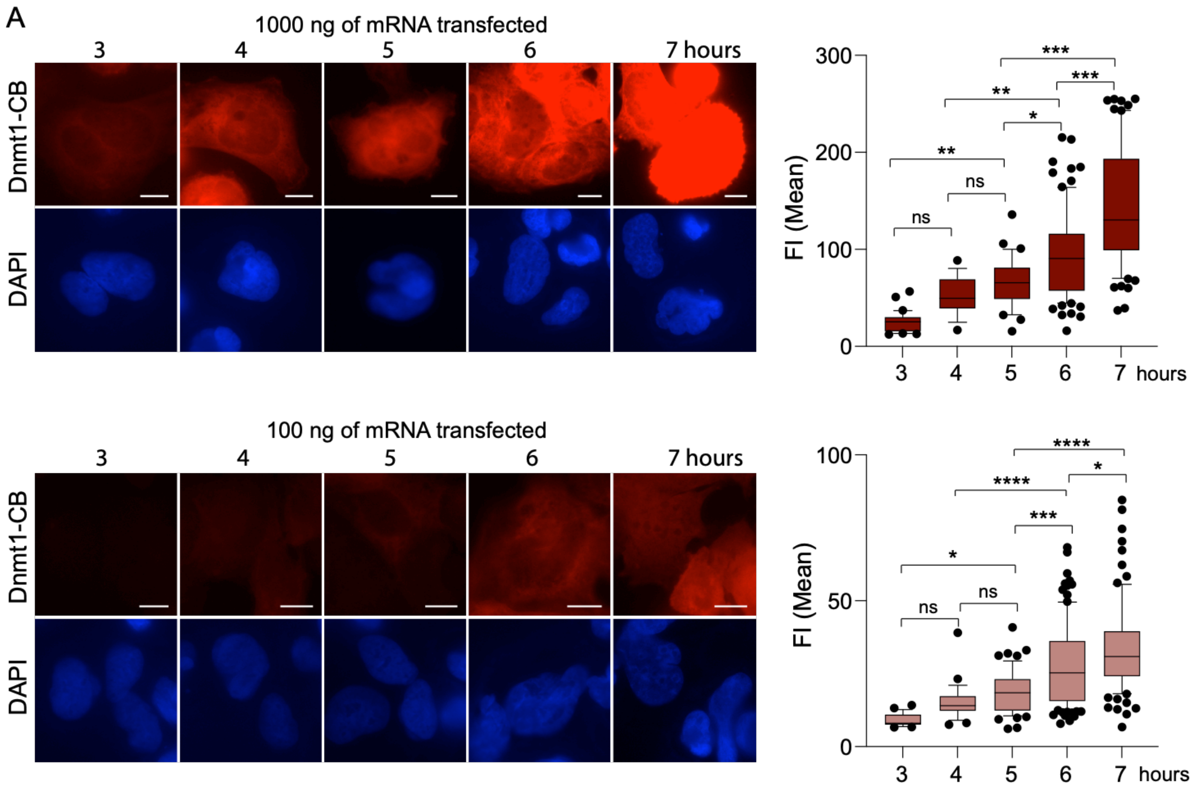


**Figure S3.** Dnmt1 chromobody-expression time points. MRC5 cells were transfected with either 1000 ng or 100 ng of mRNA encoding Dnmt1-chromobody. Cells were harvested for fluorescence imaging at different times after transfection, as indicated. Images with an identical exposure time were taken for fluorescence intensity (FI) analysis. Scale bar, 10 μm.
